# Crystal Structure of a Natural Light-Gated Anion Channelrhodopsin

**DOI:** 10.1101/405308

**Authors:** Hai Li, Chia-Ying Huang, Elena G. Govorunova, Christopher T. Schafer, Oleg A. Sineshchekov, Meitian Wang, Lei Zheng, John L. Spudich

**Affiliations:** Center for Membrane Biology, Department of Biochemistry and Molecular Biology, University of Texas Health Science Center – McGovern Medical School, Houston, TX 77030, USA; Swiss Light Source, Paul Scherrer Institute, CH-5232 Villigen PSI, Switzerland

## Abstract

The anion channelrhodopsin *Gt*ACR1 from the alga *Guillardia theta* is a potent neuron-inhibiting optogenetics tool. Presented here, its X-ray structure at 2.9 Å reveals a tunnel traversing the protein from its extracellular surface to a large cytoplasmic cavity. The tunnel is lined primarily by small polar and aliphatic residues essential for anion conductance. A disulfide-immobilized extracellular cap facilitates channel closing and the ion path is blocked mid-membrane by its photoactive retinylidene chromophore and further by a cytoplasmic side constriction. The structure also reveals a novel photoactive site configuration that maintains the retinylidene Schiff base protonated when the channel is open. These findings suggest a new channelrhodopsin mechanism, in which the Schiff base not only controls gating, but also serves as a direct mediator for anion flux.

## Introduction

Anion channelrhodopsins (ACRs) are light-gated anion channels first discovered in the cryptophyte alga *Guillardia theta* (*Gt*ACR1 and *Gt*ACR2) (*1*). Their large Cl^-^ conductance makes *Gt*ACRs and several later found ACRs (*2, 3*) the most potent neuron-silencing optogenetic tools available. *Gt*ACRs have proven to be effective inhibitors of neural processes and behavior in flies (*4-6*), worms (*7*), zebrafish (*8*), ferrets (*9*), and mice (*10-13*). The atomic structure of an ACR is essential for elucidating the mechanism of the unique natural function of light-gated anion conductance through biological membranes. Also, understanding ACR mechanisms at the atomic scale would enable rational engineering to tailor their use as optogenetic tools.

Of the 35 ACR homologs found in cryptophyte algae (*3, 14, 15*), *Gt*ACR1 is the best characterized in terms of its gating mechanism and photochemical reaction cycle (*16, 17*), and also is the only ACR for which light-gated anion conductance has been proven to be maintained *in vitro* in a purified state (*18*) further recommending it as the preferred ACR for crystallization.

The most closely related molecules to ACRs are cation channelrhodopsins (CCRs) from chlorophyte algae (*19*). The best characterized CCRs are channelrhodopsin-2 (*Cr*ChR2) (*20*), a membrane-depolarizing phototaxis receptor from *Chlamydomonas reinhardtii* (*21*), and C1C2, a chimera of *Cr*ChR2 and its paralog *Cr*ChR1 (*22*). Atomic structures of C1C2 and *Cr*ChR2 have been obtained by X-ray crystallography (*22, 23*).

The two channelrhodopsin families exhibit large differences in their sequences and photochemistry(*19*): (i) ACRs conduct only anions with complete exclusion of cations, even H^+^ for which CCRs exhibit their highest relative permeabilty; (ii) ACRs are generally more potent; e.g., *Gt*ACR1 exhibits 25-fold higher unitary conductance than *Cr*ChR2; (iii) The retinylidene Schiff base in the photoactive site deprotonates prior to channel opening in CCRs (*24*) and, in contrast, in *Gt*ACR1 remains protonated throughout the lifetime of the open-channel state with deprotonation correlated with the initial phase of channel closing (*17*).

## Overall GtACR1 structure

The *Gt*ACR1 protein was expressed in insect cells and purified as a disulfide-crosslinked homodimer (Fig. S1). We obtained lipidic cubic phase (LCP) crystals of *Gt*ACR, applied the continuous grid-scan method (*25*) to frozen LCP samples, and determined the structure at 2.9 Å resolution using molecular replacement (Fig. 1, Table S1). Each asymmetric unit contains a *Gt*ACR1 homodimer molecule (Fig. S2). Each monomer is composed of an extracellular cap domain, seven transmembrane helices (TM1-7), and a cytoplasmic loop at the carboxyl-terminus (Fig. 1). In the extracellular domain, two kinked *α*-helices from the amino-terminal fragment and a *β*-hairpin from the TM2-3 loop lay on the interface of the membrane domain. The *Gt*ACR1 homodimer is stabilized by TM3 and TM4 interactions between monomers and further by an intermolecular disulfide bridge formed by the C6 residues (Fig. 1A-B). Since TM5-7 are much longer than TM1-4, this dimeric arrangement creates a large funnel-shaped cytoplasmic cavity (∼18 Å deep and ∼28 Å wide). Despite the modest ∼24% amino acid sequence identity between *Gt*ACR1 and C1C2/*Cr*ChR2, the structure of each *Gt*ACR1 protomer can be superposed well (Fig. S3) with either of the two CCR structures (r.m.s.d. 0.9 Å) indicating that these functionally distinct channelrhodopsins share a common TM helical scaffold conformation in their closed states.

**Fig. 1.**
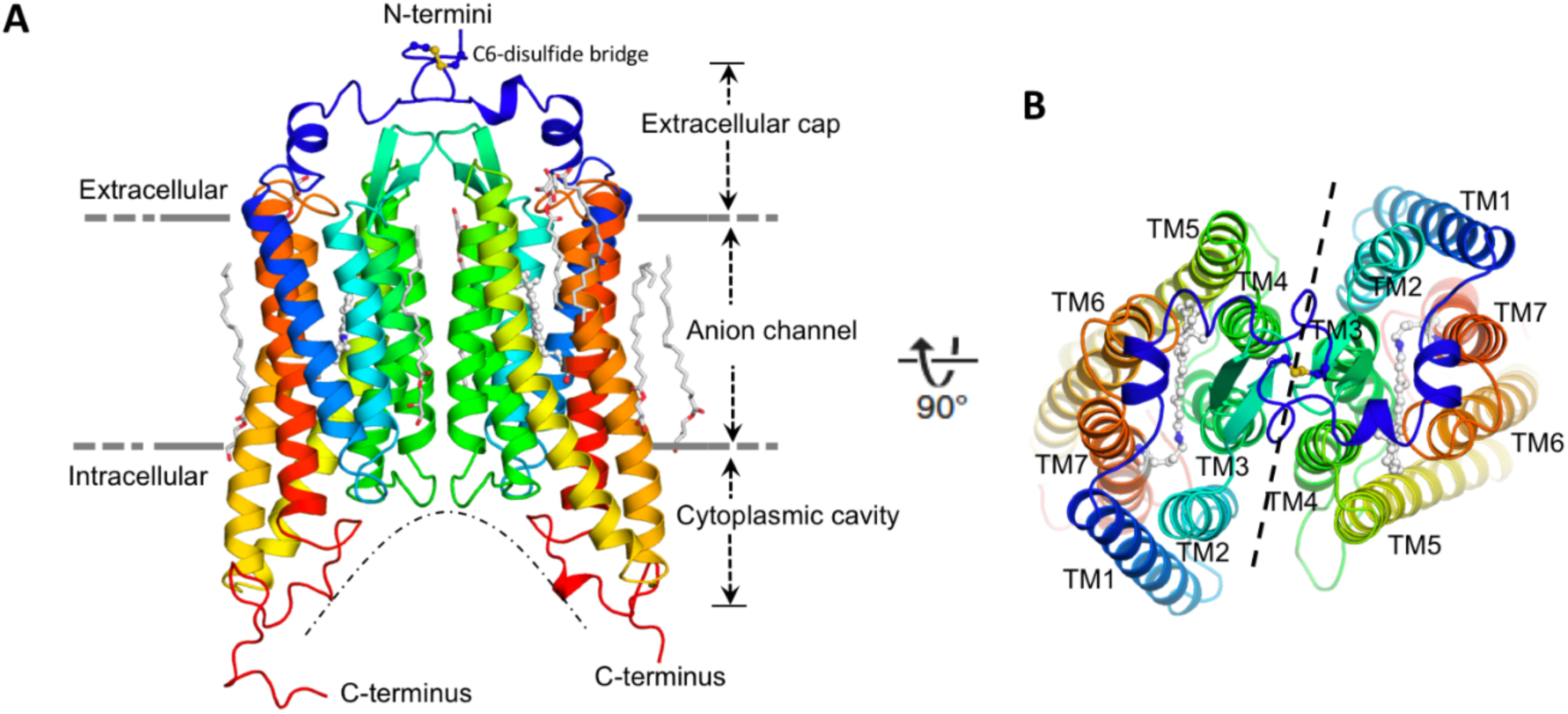
Overall structure of the *Gt*ACR1 homodimer. Side **(A)**, and top (**B**) views. Each *Gt*ACR1 protomer is depicted in cartoon with the N-termini in *blue* and the C-termini in *red*. Retinal prosthetic groups (stick-balls) are embedded in the 7TMs. The intermolecular disulfide bridge is formed by C6 (*yellow* sticks). Resolved monoolein lipids are shown as sticks.

## The anion conductance pathway

### Overview

A continuous tunnel spanning through the protein from the extracellular to cytoplasmic surface was detected in each *Gt*ACR1 protomer (Fig. 2, S4A). The tunnel, assembled by TM1-3 and 7, starts from an electropositive port on the extracellular surface, intersects the retinylidene Schiff base in the middle of the membrane, and ends at an intracellular port deeply embedded in the large dimeric cavity. In contrast, only a partial tunnel open on the extracellular side was found in C1C2 (*22*) (Fig. S4B), and no tunnel open to either surface was detected in *Cr*ChR2.

**Fig. 2.**
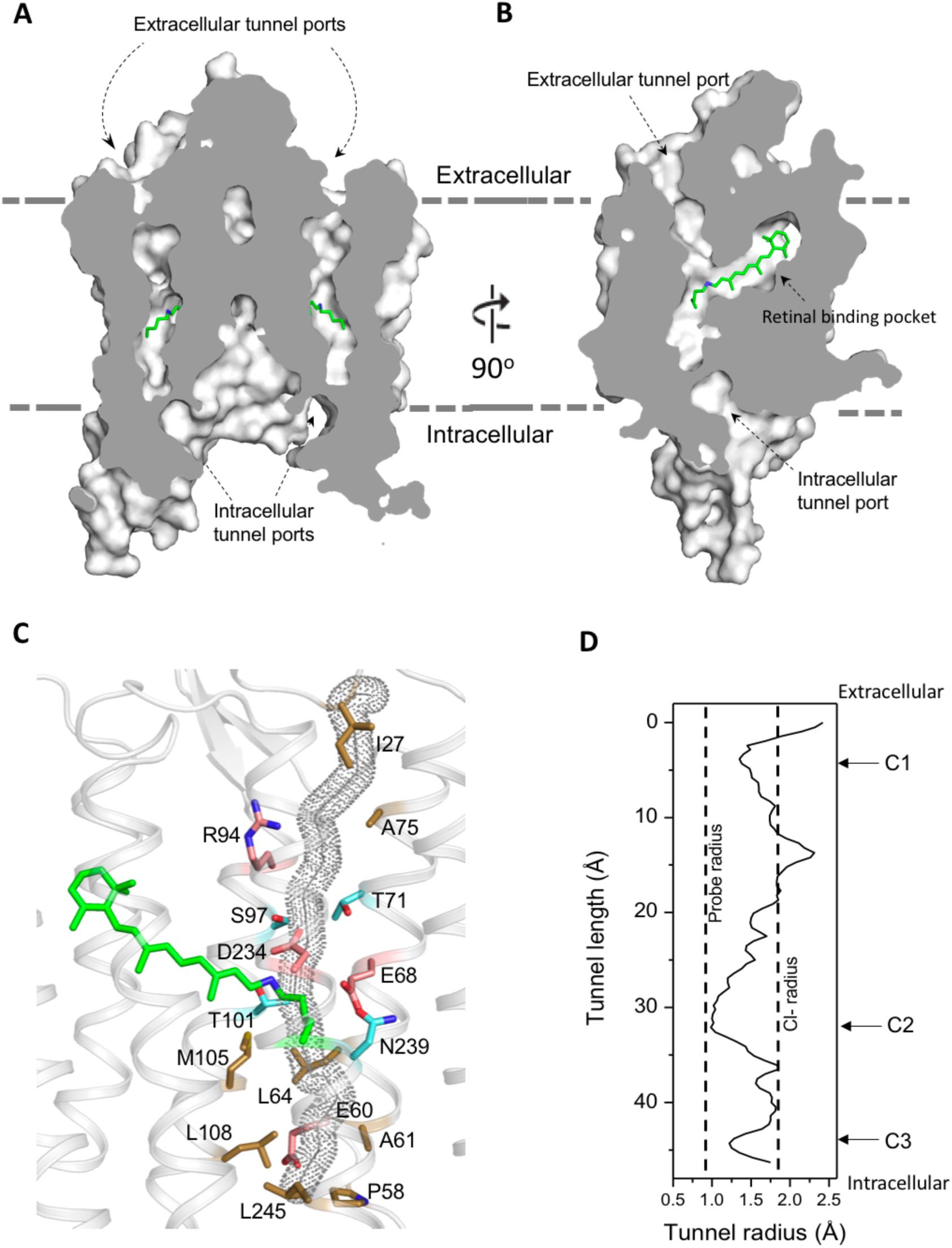
The dark state tunnel of *Gt*ACR1. A cross-section view of a *Gt*ACR1 dimer showing two continuous intramolecular tunnels traversing from extracellular ports to the cytoplasmic cavity; retinal (*green*). **(B)** A cross-section view of a *Gt*ACR1 protomer showing the conformation of the transmembrane ion tunnel and retinal binding pocket connected at the retinylidene Schiff base position. **(C)** The tunnel (*dots*) predicted by CAVER with tunnel-lining residues (*sticks*): charged (*red*), polar (*cyan*), and apolar residues (*clay*). **(D)** The tunnel profile of *Gt*ACR1 predicted by CAVER; the arrows indicate three constrictions C1-C3.

Despite the high similarity of the TM helix scaffolds of *Gt*ACR1 and C1C2/*Cr*ChR2, the tunnel of *Gt*ACR1 is primarily lined by small polar and aliphatic residues (Fig. 2C) in contrast to charged residues in the corresponding positions in C1C2 and *Cr*ChR2: A75 vs E136/E97 (C1C2/*Cr*ChR2 numbering), T71 vs K132/K93; S97 vs E162/E123, A61 vs E122/E83, and L108 vs H173/H134 (Fig. S5 top). Tunnel-lining residues also include R94 (R159/R120) and D234 (D292/D253) (Fig. S5 bottom), highly conserved in the photoactive sites of microbial rhodopsins, and E68 (E129/E90), characteristic of both ACRs and chlorophyte CCRs. The differences in *Gt*ACR1 from the CCR structures significantly reduce the negativity of the putative channel pore lining consistent with anion vs cation conductance.

### The extracellular port cap

A unique structural feature is found in the extracellular domain of *Gt*ACR1. In addition to the disulfide link between the two protomers, an intraprotomer disulfide bridge is formed between C21 from the amino-terminal segment and C219 within the TM6-7 loop (Fig. 3A). This intramolecular crosslink immobilizes the kinked helices to the retinal-conjugated TM7, and encaps a hydrophobic part of the segment on the extracellular tunnel entry (Fig. 3B). Disrupting this extracellular loop conformation, either by truncation of the amino-terminal loop (Δ1-25) or by substituting C21 and C219 with serine to abolish the intramolecular disulfide, resulted in slowed channel closing (Fig. 3C). Both C21 and C219 are highly conserved in ACRs (*2*), but not in CCRs, revealing a role of this intramolecular disulfide bridge specific to the ACR family.

**Fig. 3.**
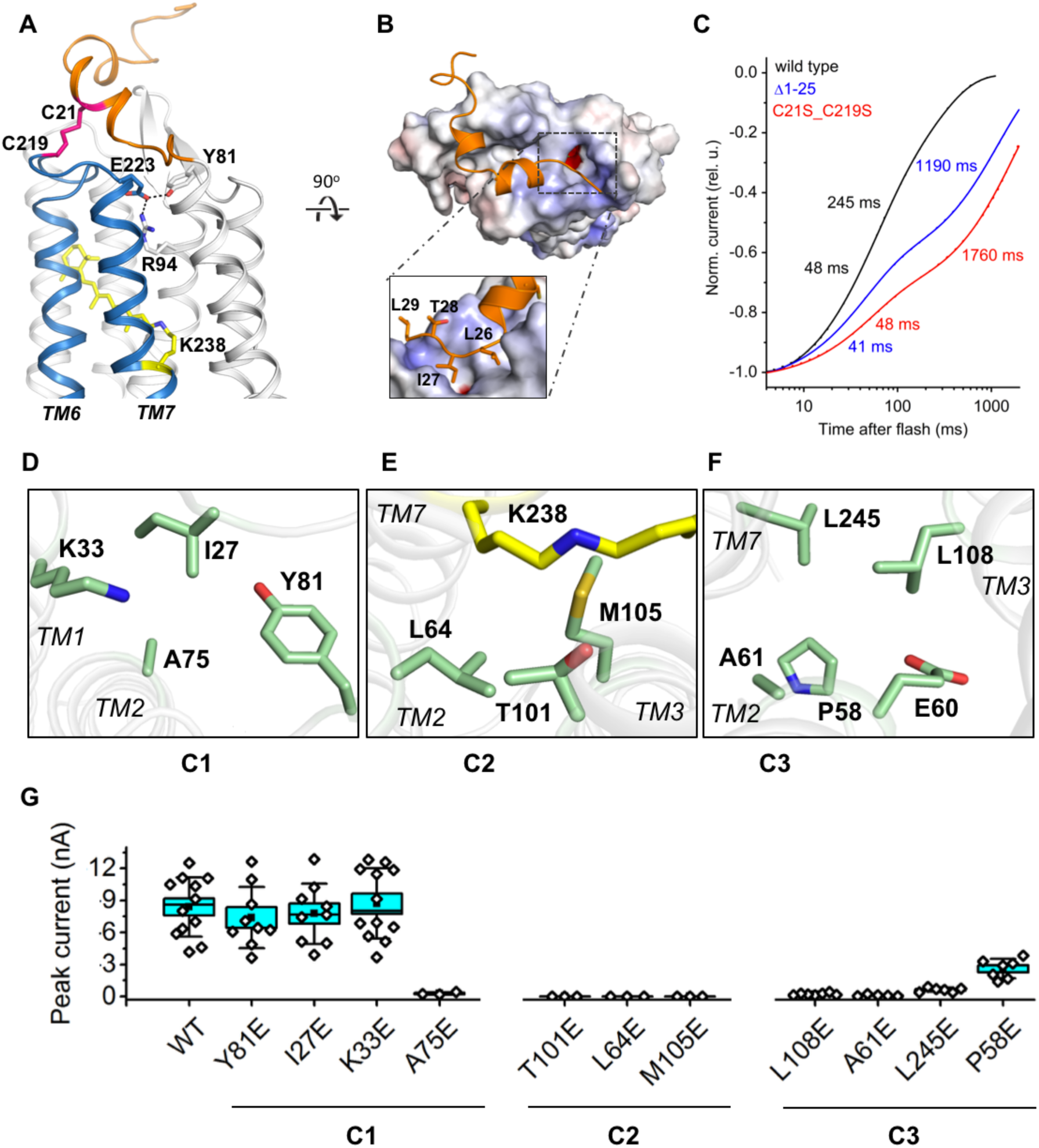
Features of the ion pathway of *Gt*ACR1. **(A)** The extracellular loop (*orange*) immobilized with retinal (*yellow*)-conjugated TM6-7 (*blue*) by an intracellular C21-C219 disulfide bridge (*red*); an H-bond network (*black dashed lines*) formed by residues (*sticks*) near the extracellular port. **(B)** The hydrophobic segment (*orange*) blocks the extracellular port rendered by the electrostatic potential surface calculated by APBS. Rectangle: closer view of the peptide cap conformation. **(C)** Decay kinetics of laser flash-evoked photocurrents by the wild-type *Gt*ACR1 and indicated mutants. **(D-F)** The structure of the three constrictions: C1 (D), C2 (E), and C3 (F). **(G)** Peak photocurrents generated by Glu mutants of the constriction residues.

### Ion pathway constrictions

The intramolecular tunnel in *Gt*ACR1, presumably indicating the anion conductance pathway, is constricted at three positions as detected using the program CAVER (probe radius 0.9 Å) (*26*): at the extracellular port (C1), near the photoactive retinylidene Schiff base (C2), and at the cytoplasmic side (C3) (Fig. 3D-F). Near the extracellular port, the C1 constriction (Fig. 3D) is stabilized by an H-bond network adjacent to the disulfide-immobilized extracellular cap and formed by the side chains of Y81, R94 and E223 (Fig. 3A). The mutation R94A nearly abolished Cl^-^ conductance (Fig. 4D), whereas the mutation E223Q resulted in ∼10-fold slowing of the current decay rate (*16*), similar to that observed in the C21S_C219S mutant. These results suggest that the combination of the H-bond network of E223 and its neighbouring intraprotomer disulfide bridge controls the rate of channel closing in the extracellular region and stabilize the essential residue R94 in the closed state.

**Fig. 4.**
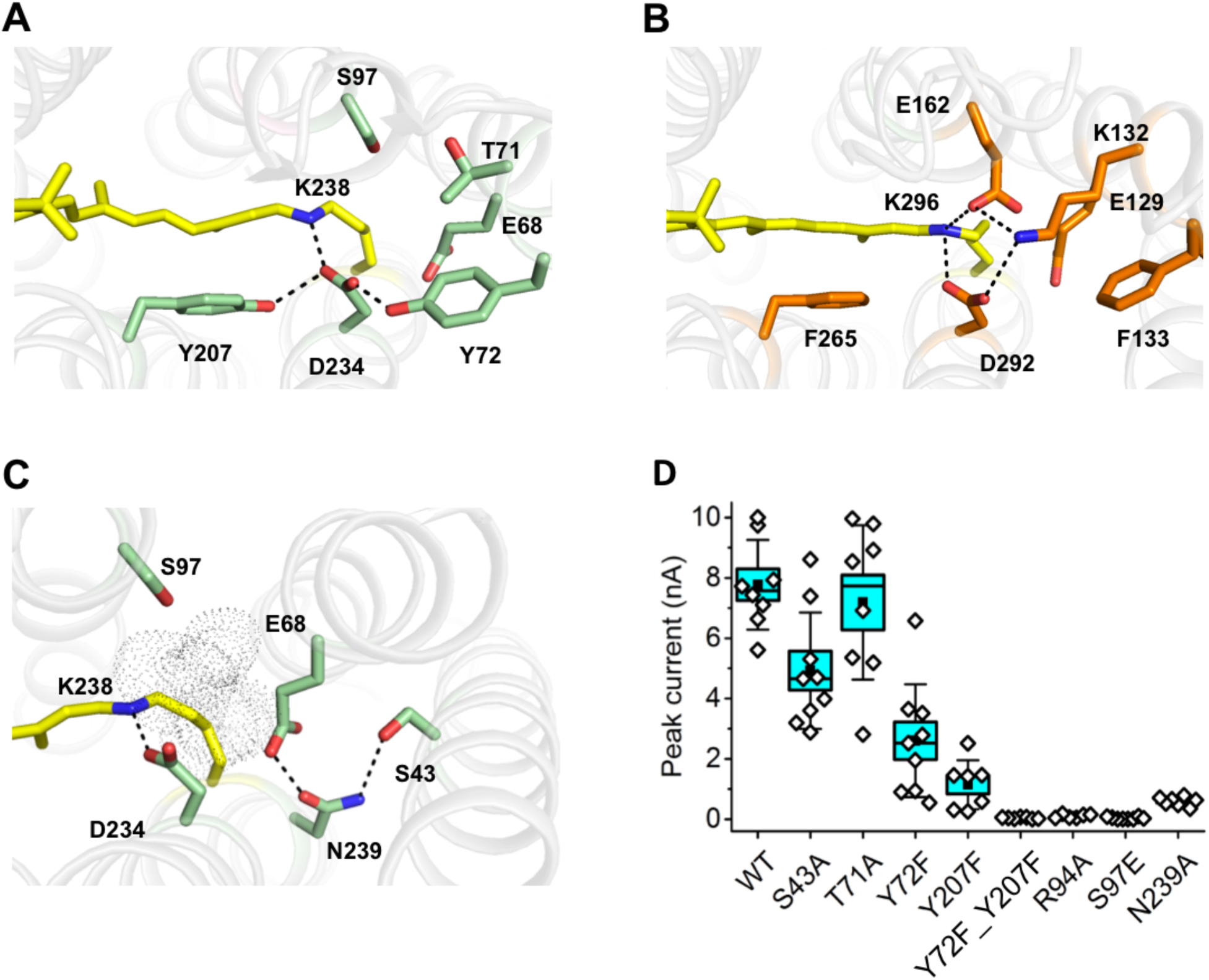
Conformation of the Schiff base region of *Gt*ACR1. **(A-B)** Structural comparison shows different H-bond networks (dashed lines) in *Gt*ACR1 **(A)** and C1C2 **(B)**. **(C)** the H-bond network in the ENS triad of *Gt*ACR1. The tunnel (*pink* surface) was detected by CAVER. **(D)** Peak photocurrents generated by the wild-type *Gt*ACR1 and indicated mutants.

The narrowest constriction C2 lies at the photoactive site and is formed by the side chains of T101, L64, and M105 (Fig. 3E). Four of the five residues that form the intracellular constriction C3 (L108, A61, E60, L245 and P58) (Fig. 3F) are in corresponding positions as the residues that form the “intracellular gate” in CCRs (*27*), but in *Gt*ACR1 and other ACRs only E60 (E121/E82) is shared with CCRs. The *Gt*ACR1 structure that we obtained from dark-grown crystals is presumably the dark (closed) state of the channel. To examine the role of these contriction-forming residues in the channel open state, we scanned the tunnel constrictions with Glu substitutions and measured photocurrents in the respective mutants. While effects of most mutations (except A75E) at the C1 position were negligible, perturbation of any residues at C2 or C3 greatly reduced or eliminated the photocurrents (Fig. 3G), suggesting that in the open conformation the channel is wider in the extracellular portion and more narrow in its central and intracellular stretches.

### The retinal polyene chain and ring

In the middle of the protein, all-*trans*-retinal covalently bound by a Schiff base linkage to K238 is found in an elongated cavity formed by conserved hydrophobic residues. While the conformations of the retinal polyene chain are nearly identical in *Gt*ACR1 and C1C2/*Cr*ChR2, the presence of F160 in *Gt*ACR1 (G224/G185 in C1C2/CrChR2, respectively) pushes the *β*-ionone ring towards the extracellular side by 1.2 Å (Fig. S6).

### The retinylidene Schiff base region

Remarkable structural differences between *Gt*ACR1 and the two CCRs are found in the retinylidene Schiff base environment. In C1C2 and *Cr*ChR2 the protonated Schiff base participates in a quadruple salt-bridge network formed with D292/D253, E162/E123 and K132/K93 sidechains (Fig. 4B). However, this strong network is absent in the *Gt*ACR1 structure due to the replacement of E162/E123 and K132/K93 with smaller non-carboxylate residues S97 and T71, respectively (Fig. 4A). D234 is the only residue directly interacting with the protonated Schiff base in *Gt*ACR1, and its electrostatic interaction is weakened by two H-bonds from tyrosine residues Y72 and Y207 (Fig. 4A). The proton pump bacteriorhodopsin exhibits similar tyrosinyl H-bond-weakened interactions of D212, the residue in the corresponding position as D234. The interactions prevent D212 from accepting the Schiff base proton, which is transferred instead to D85 in the proton release pathway (*28*). Resonance Raman and UV-vis absorption spectra of the D234N mutant of *Gt*ACR1 indicate that D234 is similarly neutral and not a Schiff base proton acceptor (*17*) (*29*). The dark structure therefore appears to explain the persistence of a protonated Schiff base throughout the lifetime of the open channel conformation in *Gt*ACR1.

In *Cr*ChR2, photoisomerization of the Schiff base rapidly disrupts the strong salt-bridged network by inducing transfer of the Schiff base proton to D253 or E123 in ∼10 μs, prior to channel opening (*24*). In contrast, the *Gt*ACR1 Schiff base remains protonated throughout the lifetime of the open channel conformation in *Gt*ACR1 and deprotonation of the Schiff base proton occurs late in the photocycle (∼20 ms) correlated with fast channel closing (*17*). Unlike in the salt-bridge network around the Schiff base in the CCRs (Fig. 4B), in *Gt*ACR1 no immediate proton accepting residue is available in the vicinity of the protonated Schiff base and therefore later structural changes are required to enable Schiff base proton transfer, possibly to E68 (Fig. 4A).

The protonated Schiff base, centered along the anion path in ACRs, may play a direct role in anion conduction. Supporting this possibility, late deprotonation of the Schiff base after channel opening occurs in all three ACRs so far examined: *Gt*ACR1 and *Gt*ACR2 (*17*) and *Psu*ACR1 (*15*), yet Schiff base deprotonation after channel opening is not known to occur in any CCR. Also supporting a functional role, the mutant S97E, in which a potential Schiff base proton acceptor is placed at the corresponding position in *Gt*ACR1 as in CCRs and many other microbial rhodopsins, results in (i) appearance of fast Schiff base deprotonation, and (ii) a >30-fold suppression of the amplitude of the chloride photocurrent in the dark adapted state (*17*). Furthermore, the double mutation Y207F/Y72F, expected to release inhibition of D234 as a proton acceptor, decreased the photocurrent amplitude to negligible (Fig. 4D).

### The ENS triad

E68, a glutamyl residue near the Schiff base constriction in the channel, forms an H-bond network with N239 and S43 (Fig. 4C) with a geometry similar to that of a homologous triad (E129/E90, N297/N258, and S102/S63) referred to as “the central gate” in C1C2 and *Cr*ChR2. In CCRs, the triad blocks the cation path from the extracellular bulk phase (*27*) and the glutamyl residue contributes to cation selectivity over anions (*30*). In contrast, in *Gt*ACR1 the ENS triad does not occlude the tunnel (Fig. 4C), but E68 is functionally important in channel gating and may serve as a Schiff base proton acceptor (*16*). The three residues in the ENS triad appear to have distinct roles; i.e. the substitution S43A had little effect on Cl-conductance, whereas the mutation N239A nearly eliminated the photocurrent (Fig. 4D). Given its location between C2 and C3, N239 may assist moving anions between the Schiff base and the cytoplasmic port (Fig. 2C). Additionally, the distribution of apolar residues in this portion of the channel would also facilitate quick movements of anions as has been proposed for the CLC channel (*31*).

Despite the large phylogenetic difference between cryptophyte ACRs and chlorophyte CCRs, their helical scaffolds are little changed. However, the *Gt*ACR1 structure reveals fundamentally different chemistry built within their common scaffold. The preexisting full-length tunnel, the location of the retinylidene photoactive site directly in the ion path, the maintenance of a net positive charge on the site’s Schiff base, and the novel extracellular cap, provide a first view of the structural basis of light-gated anion conductance.

## Acknowledgements

This work was supported by National Institutes of Health Grants R01GM027750 and U01MH109146, the Hermann Eye Fund, and Endowed Chair AU-0009 from the Robert A. Welch Foundation to J.L.S. C.-Y.H. was partially supported by the European Union’s Horizon 2020 research and innovation programme under the Marie-Skłodowska-Curie grant agreement No. 701647. Author contributions: J.L.S. and L.Z. designed the research; all authors performed research and/or analyzed results and all contributed to the manuscript. Atomic coordinates and structure factors for the reported crystal structure have been deposited with the Protein Data Bank (PDB) under the accession code 6EDQ.

